# Dynamics of interactions between three major respiratory pathogens in reconstituted human epithelium

**DOI:** 10.1101/2025.11.17.688795

**Authors:** Aurélien Gibeaud, Charles Terra, Émilie Portier, Pascal Lukas, Audrey Le Gouellec, Jeremie Guedj, Frederik Graw, Andrés Pizzorno, Olivier Terrier

## Abstract

Respiratory viral coinfections pose a substantial global health burden, yet the underlying virus–virus interactions remain incompletely understood. Here, we systematically examined the interplay among influenza A virus (IAV), SARS-CoV-2, and respiratory syncytial virus (RSV) using a reconstituted human airway epithelium model. We monitored viral replication dynamics and host transcriptional responses under both simultaneous and sequential infection conditions. Our results reveal complex, asymmetric interactions strongly influenced by infection timing and sequence. IAV exerted a pronounced inhibitory effect on SARS-CoV-2, mediated mainly by a robust type III interferon response. However, transient early enhancement of SARS-CoV-2 by IAV and subsequent bidirectional inhibition were also observed. Transcriptomic profiling identified coinfection-specific gene expression signatures enriched for metabolic and cell death pathways. In IAV–RSV coinfections, IAV generally suppressed RSV replication; strikingly, prior RSV infection significantly enhanced subsequent IAV replication, potentially through RSV-induced cellular remodeling or syncytia formation. Overall, host responses to coinfection were integrated and non-additive, with distinct transcriptional programs suggesting activation of unique cellular pathways beyond canonical antiviral responses. Together, these findings highlight the pivotal role of innate immunity and the order of infection in determining the outcome of respiratory viral coinfections, providing mechanistic insights with implications for clinical management and epidemic modeling.

**Author Summary:** Respiratory viruses often circulate simultaneously; however, their interactions within the same host remain poorly understood. We used a reconstituted human airway epithelium model to examine coinfections involving influenza A virus (IAV), SARS-CoV-2, and respiratory syncytial virus (RSV). Our findings reveal that these viruses do not simply compete or coexist: their interactions are dynamic, asymmetric, and highly dependent on the order in which infections occur. IAV generally suppresses SARS-CoV-2 replication through a strong type III interferon response, although brief early enhancement and later mutual inhibition can occur. Transcriptomic profiling reveals that coinfection triggers a complex array of host responses, including alterations in metabolic and cell-death pathways. In IAV/RSV coinfections, IAV consistently inhibits RSV; however, prior RSV infection markedly boosts subsequent IAV replication—likely due to RSV-induced cellular remodeling, such as syncytia formation. These transcriptional signatures point to altered cell adhesion and motility as key features of such interactions. Overall, our results demonstrate that innate immunity and the sequence of infection shape coinfection outcomes in non-additive ways, driving distinct cellular responses beyond classical antiviral pathways. Understanding these mechanisms is essential for improving clinical management and guiding the development of targeted antiviral strategies.

## INTRODUCTION

Respiratory viral infections represent a significant global health burden, ranking among the leading causes of illness and mortality worldwide [1,2]. In 2019, approximately 17.2 billion cases of upper respiratory infections were estimated, constituting over 40% of all illnesses globally [3]. The respiratory viruses responsible for these infections notably include respiratory syncytial virus (RSV) and influenza A viruses (IAV), which collectively cause a broad spectrum of clinical manifestations ranging from mild upper respiratory symptoms to severe lower respiratory tract disease [4]. The transition of SARS-CoV-2 from pandemic to endemic status has introduced an even more complex landscape of viral co-circulation, reinforcing interest in how these viruses interact at different scales. Indeed, respiratory viruses can infect the same host simultaneously or sequentially, leading to coinfections. The increasing use of multiplex diagnostic panels has revealed that viral codetection is more common than previously appreciated, with coinfections reported in 3% to 26% of hospitalized patients with respiratory infections [5]. Coinfections can involve homologous viruses (from the same family), heterotypic viruses (different strains or subtypes within a species), or heterologous viruses (from different families), underscoring the diverse possible interaction scenarios [6].

The clinical implications of respiratory viral coinfections remain an area of active investigation and debate. Coinfections are particularly prevalent in vulnerable populations such as children, the elderly, and immunocompromised individuals, raising concerns about potential impacts on disease severity, hospitalization risk, and therapeutic management [7,8]. Of particular interest are coinfections involving SARS-CoV-2 and IAV, as well as IAV and RSV, given their significant contributions to respiratory morbidity and mortality. Since the onset of the COVID-19 pandemic in 2020, the interplay between SARS-CoV-2 and other respiratory viruses has attracted attention. Several studies have shown that more than 10% of SARS-CoV-2 patients were co-infected with other respiratory viruses [9,10]. On the other hand, epidemiological studies indicate that individuals infected with influenza have a substantially reduced risk of testing positive for SARS-CoV-2, suggesting potential viral interference and/or more complex interactions [7]. However, when coinfections do occur, they are associated with increased disease severity and higher mortality rates than mono-infections, highlighting the need for vigilant clinical management [7]. Similarly, interactions between influenza and RSV have been documented, with evidence of competition that may influence epidemic timing and clinical outcomes [11,12]. These clinical challenges underscore the importance of understanding the epidemiology, severity, and optimal management strategies for viral coinfections.

The underlying mechanisms governing viral coinfections are complex and extend beyond the classical notion of viral interference, which historically described negative interactions in which infection by one virus inhibits a second virus, often mediated by the host interferon (IFN) response [13]. Viruses can either compete for resources and sustain a negative interaction or on the contrary, develop a positive interaction by enhancing each other’s ability to drive their replication cycle [6]. Current evidence indicates that the nature of virus-virus interactions depends on multiple factors, including the identities and families of the viruses involved, the sequence and timing of infections, and the dynamics of the host immune response. For example, *in vitro* studies have demonstrated that prior infection with human rhinovirus or IAV can block SARS-CoV-2 replication in respiratory epithelial cells by inducing an IFN response, providing a clear case of viral interference mediated by innate immunity [14,15]. Similar observations were made for the coinfection between IAV and RSV [16–19]. For example, Gilbert-Girard and colleagues demonstrated that IAV interferes with RSV replication *in vivo*; however, the opposite was not observed. In reconstituted human airway epithelia, viral interference was dependent on the timing and sequence of infections but not on differential interferon susceptibilities [16]. Conversely, certain studies have suggested positive interactions, such as hybrid viral particles formed between influenza A virus (IAV) and RSV that broaden receptor tropism and may enhance infectivity [20].

The impact of infection order is also critical; we have previously shown that the outcome of SARS-CoV-2 coinfections with IAV, RSV, human metapneumovirus (hMPV), or rhinovirus (hRV) in reconstituted human airway epithelia varied significantly depending on the nature of the virus and the sequence of infection, affecting viral replication kinetics and innate immune responses [21]. Moreover, the antiviral IFN response, particularly type I and III IFNs, plays a pivotal role in mediating these interactions, but IFN-independent mechanisms, such as competition for cellular receptors and resources, also contribute [6]. These findings underscore the need for more systematic and multifaceted experimental approaches, incorporating relevant human airway models and considering factors such as timing, viral strain, and host factors, to fully elucidate the mechanisms of respiratory viral coinfections.

In this study, we aimed to explore interactions among IAV, SARS-CoV-2, and RSV across different coinfection scenarios in reconstituted human airway epithelia by monitoring and comparing multiple viral and host parameters during the course of infection. Our results confirm that the nature of interactions between viruses depends heavily on the timing and sequence of infection. The nature of the primary infection and the associated host response, including the innate immune response, are key factors that determine the outcome of the coinfection. Furthermore, beyond mutual negative interactions, we have identified heterogeneous interactions (positive/negative) in certain cases, reflecting an even greater degree of complexity.

## RESULTS

### Simultaneous infection of HAE by IAV and SARS-CoV-2

We first focused our investigation on the interactions between IAV and SARS-CoV-2 by performing infection kinetics in reconstituted human airway epithelia derived from multiple donors (n = 10), under various coinfection scenarios. We monitored viral replication dynamics (genome copy numbers) together with several host parameters, including epithelial integrity (measured by transepithelial electrical resistance, TEER) and the secretion of IL-6 and IFN-λ1, respective hallmarks of inflammatory and antiviral immune responses (**Fig. 1A-C**). The multiplicity of infection (MOI) was set to 0.1 for IAV and 0.01 for SARS-CoV-2 in order to achieve comparable replication kinetics over a 72-hour post-infection (hpi) period (**Fig. 1A**). During single IAV infection, viral genome copy numbers increased over time, peaking at 48 hpi and slightly declining at 72 hpi (blue bars, **Fig. 1B**). This replication dynamic was associated with a sharp decrease in TEER, reflecting severe disruption of epithelial integrity (blue bars, **Fig. 1D**), as well as a strong induction of IL-6 secretion starting at 48 hpi and of IFN-λ1 secretion much earlier, from 10 hpi onwards (blue bars, **Fig. 1E** and **1F**). In single SARS-CoV-2 infection, genome copy numbers also progressively increased (yellow bars, **Fig. 1C**), accompanied by a marked drop in TEER from 48 hpi (yellow bars, **Fig. 1D**). Infection triggered IL-6 secretion, which remained constant throughout the infection course, whereas IFN-λ1 secretion gradually increased over time, although at lower levels than those observed in IAV infection (yellow bars, **Fig. 1E** and **1F**). In the case of simultaneous infection with IAV and SARS-CoV-2, we observed a complex interplay between the two viruses. IAV replication was transiently enhanced by approximately 150% at 24 hpi, followed by a 73% reduction at 48 hpi relative to single infection (green bars, **Fig. 1B**). Conversely, SARS-CoV-2 replication was strongly inhibited, with a nearly 99% decrease in viral genome copies from 48 hpi compared with single infection (green bars, **Fig. 1C**). These findings were corroborated by confocal fluorescence microscopy at 48 hpi, which showed reduced detection of viral nucleoproteins (NP for IAV and N for SARS-CoV-2) in the coinfection condition compared with single infections, with limited overlap between the two viral signals (**Fig. 2**). Host response monitoring revealed a pronounced drop in TEER starting at 48 hpi and a progressive increase in IL-6 and IFN-λ1 secretion during simultaneous coinfection, following a pattern similar to that observed in single IAV infection (green bars, **Fig. 1D** and **1F**). We further analyzed how viral and host parameters evolved relative to one another by assessing their correlations under each experimental condition. While viral genome copy numbers for IAV and SARS-CoV-2 (GCi and GCs, **Fig. 1G**) were strongly correlated with TEER and IFN-λ1 levels during single infections, this correlation was lost for SARS-CoV-2 genome copies under simultaneous coinfection (**Fig. 1G**).

**Figure 1.**
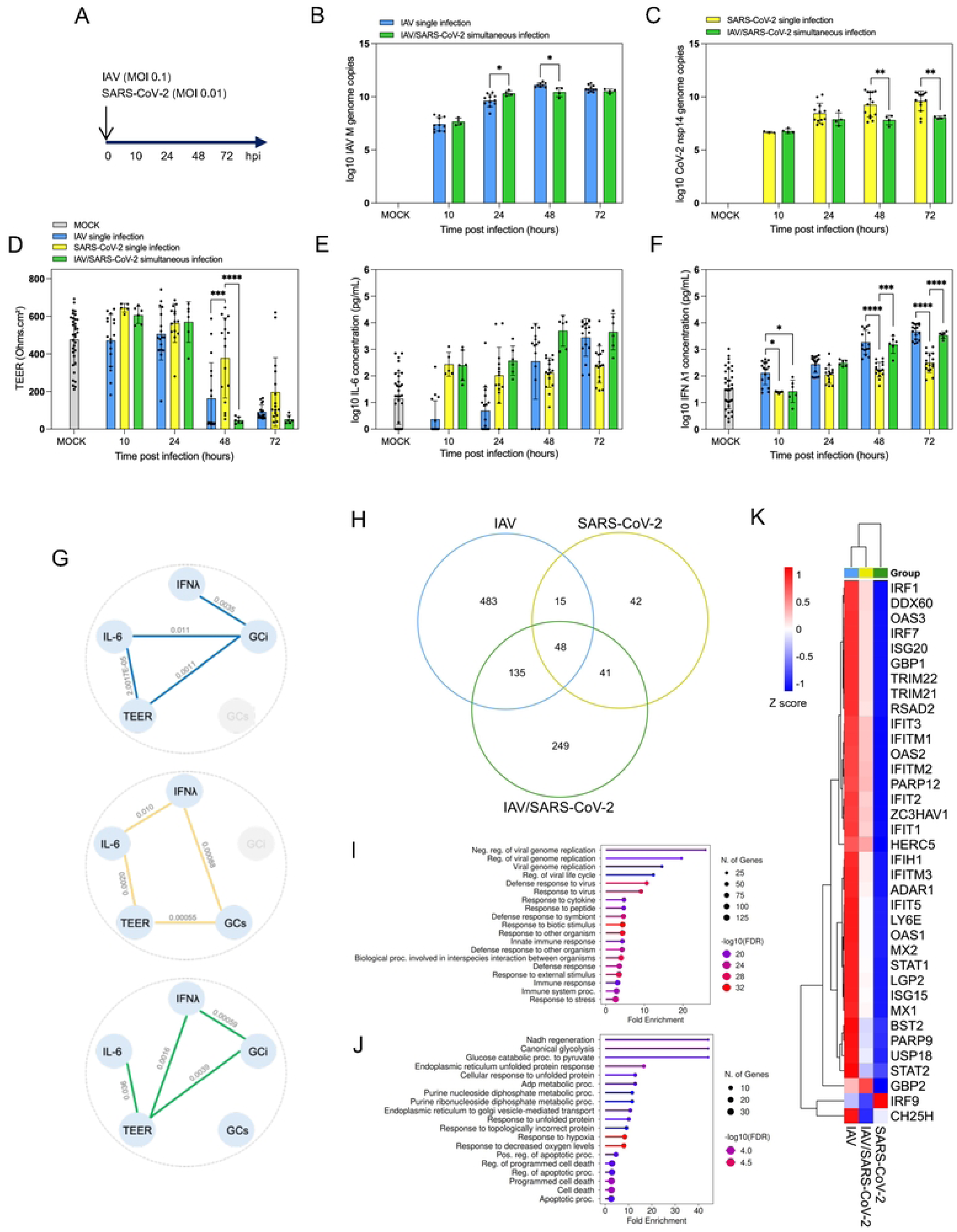
Simultaneous coinfection of IAV and SARS-CoV-2. (**A**) Reconstituted human bronchial epithelial tissues derived from six independent donors were infected or coinfected with IAV and/or SARS-CoV-2 at multiplicities of infection (MOIs) of 0.1 and 0.01, respectively. Viral replication was monitored over time by quantifying viral genome copy numbers of the IAV *M* gene (**B**) and the SARS-CoV-2 *nsp14* gene (**C**) using RT-qPCR. In parallel, several host parameters were measured, including (**D**) transepithelial electrical resistance (TEER, Ω·cm²) as a proxy for epithelial integrity, and (**E–F**) secretion of IL-6 and IFN-λ1 by ELISA to monitor inflammatory and antiviral responses. (**G**) Partial correlation analysis was performed between viral and host parameters. To further characterize host transcriptional responses to IAV and SARS-CoV-2 interaction, transcriptomic profiling was performed at 48 hpi using 3′ RNA-seq (**H–K**), selecting for each condition one representative infection from a single donor. (**H**) Differential expression analysis identified deregulated genes (DEGs) that were shared or specific among experimental conditions. Functional enrichment analysis of Gene Ontology (GO) terms (Biological Processes) was performed using either (**I**) the global or (**J**) the coinfection-specific upregulated gene signatures. (**K**) Expression profiles of 36 interferon-stimulated genes (ISGs) known to participate in IAV and SARS-CoV-2 infection were examined, representing major innate immune pathways. Data are shown as heatmaps and expressed as Z-scores. Statistical analyses for panels (**B–F**) were performed using one-way ANOVA followed by Dunnett’s, Sidak’s, and Tukey’s multiple-comparison tests. *, **, ***, and **** denote adjusted p-values of 0.05, 0.01, 0.001, and 0.0001, respectively.

**Figure 2.**
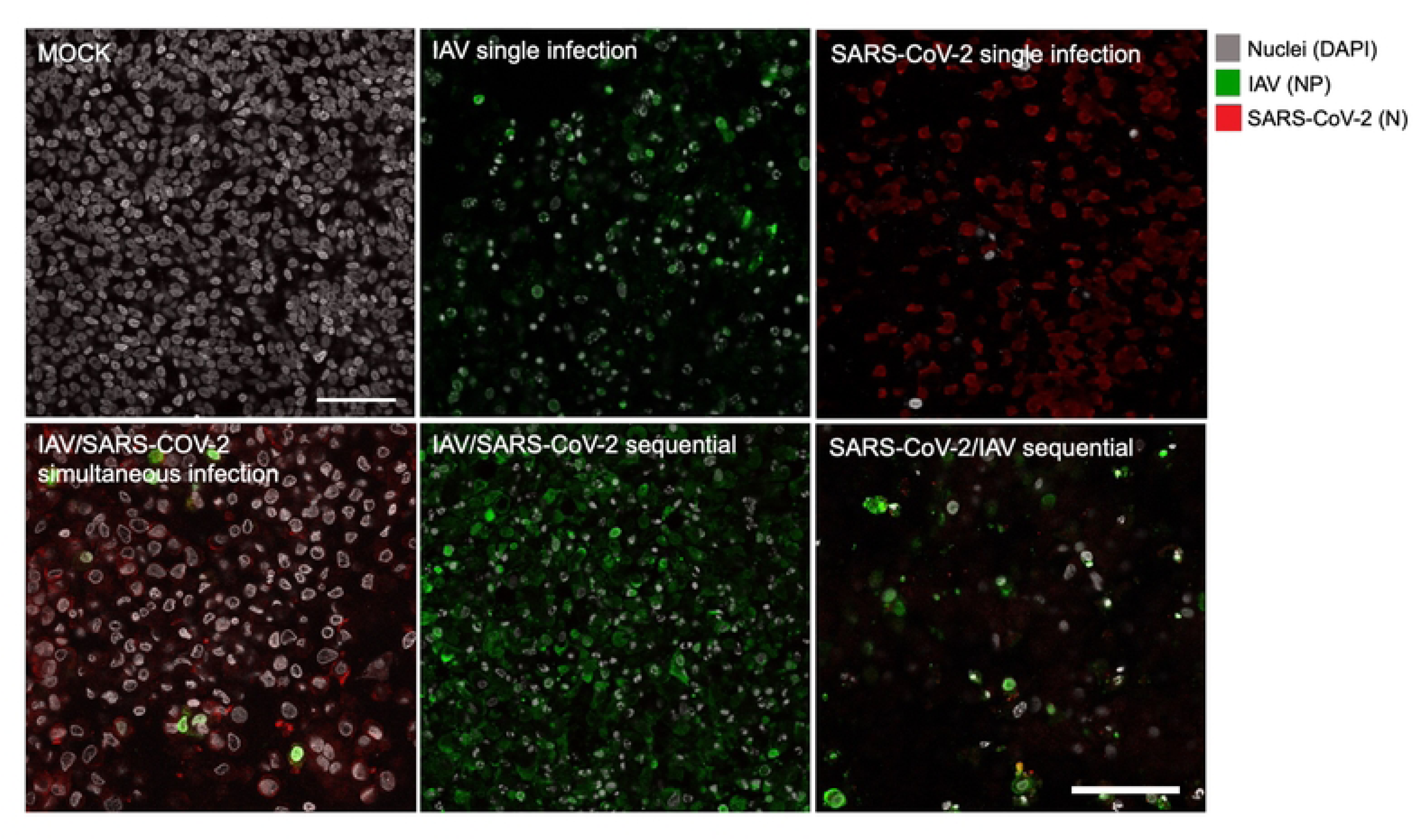
Confocal microscopy analysis of IAV and SARS-CoV-2 coinfection scenarios. Representative confocal images of single infections or coinfections (simultaneous or sequential) were acquired at 48 hours post-infection (hpi) or 48 hours post-coinfection (hpci), respectively. Confocal imaging was performed in oil immersion using a Leica SP5X microscope equipped with a 63× objective lens, and the images were processed with ImageJ Fiji software. Nuclei are shown in gray, IAV nucleoprotein (NP) in green, and SARS-CoV-2 nucleocapsid (N) in red. Scale bar: 50 µm.

To better characterize the host response during IAV/SARS-CoV-2 interaction, we performed transcriptomic profiling at 48 hpi using 3′ RNA-seq, selecting, for each experimental condition, a representative infection from a single donor. Differential gene expression analysis relative to the MOCK control (log₂ FC <-1.5 or ≥ 1.5, adjusted p-value ≤ 0.05) revealed a distinct transcriptomic signature for simultaneous IAV/SARS-CoV-2 coinfection, comprising 473 differentially expressed genes (DEGs). This signature showed 38.6 % and 18.8 % overlap with those from IAV and SARS-CoV-2 single infections, respectively (**Fig. 1H**). Notably, a large fraction of DEGs (n = 249, 52,6 %) was specific to simultaneous coinfection (**Fig. 1H**). Functional enrichment analysis of upregulated genes within the IAV/SARS-CoV-2 signature revealed significant overrepresentation of Gene Ontology (GO) terms associated with antiviral and, more broadly, immune responses biological processes (**Fig. 1I**). These enrichments were similar to those identified in single IAV and SARS-CoV-2 infections (**S1 Fig. A**) and were largely driven by DEGs common to all three signatures (**Fig. 1H**). Focusing on genes specifically upregulated during IAV/SARS-CoV-2 coinfection, functional analysis highlighted enrichment in GO terms related to cellular metabolism and cell death (**Fig. 1J**), suggesting additional pathways involved in the epithelial response in this context. We next examined the expression profiles of 36 interferon-stimulated genes (ISGs) known to participate in IAV and SARS-CoV-2 infection, representing major innate immune pathways (**Fig. 1K**). Most ISGs were robustly upregulated across all infection conditions, with a general pattern of strong induction during IAV infection, weaker induction during SARS-CoV-2 infection, and intermediate levels during coinfection (**Fig. 1K**). This trend paralleled the secretion patterns of IL-6 and IFN-λ (**Fig. 1E and 1F).** Interestingly, IRF9 expression followed an opposite trend compared to other ISGs (**Fig. 1K**).

Together, these results indicate a heterogeneous interaction at 24 hpi — positive for IAV but negative for SARS-CoV-2 — that evolves into a mutual negative interaction from 48 hpi onward. Under these experimental conditions, although both viruses exert a reciprocal inhibitory effect, the impact of IAV on SARS-CoV-2 is far more pronounced, likely due to the stronger host immune and inflammatory responses elicited by IAV. This is consistent with the higher secretion of IFN-λ1 and IL-6 (**Fig. 1E** and **1F**) and with the extensive upregulation of genes involved in these pathways by IAV compared with SARS-CoV-2 (**Fig. 1K**).

### Sequential infections between IAV and SARS-CoV-2

We next investigated sequential coinfection scenarios between IAV and SARS-CoV-2, introducing a 24-hour interval between the two infections (**Fig. 3A** and **3F**). When HAE were first infected with IAV followed by SARS-CoV-2, we observed a 283% increase in IAV genome copy numbers, compared to single IAV infection at 48 hours post-coinfection (hpci) (green bar, **Fig. 3B**). In parallel, SARS-CoV-2 replication was almost completely inhibited, showing a 99% reduction in genome copy numbers as early as 24 hpci (green bars, **Fig. 3C**). This antagonistic interaction was also confirmed by confocal fluorescence microscopy, which revealed drastically reduced SARS-CoV-2 N-protein signal in co-infected samples (**Fig. 2**). In this sequential scenario (IAV followed by SARS-CoV-2), the effect on epithelial integrity was delayed compared with single IAV infection, as indicated by TEER measurements, while the kinetics of IL-6 and IFN-λ1 secretion mirrored those observed during IAV infection alone (green bars, **Fig. 3D**). Correlation analysis among viral and host parameters revealed a strong correlation between GCi and GCs, together with a loss of correlation between GCi and TEER, consistent with the observed phenotypes (**Fig. 3E**). Conversely, in the inverse sequential scenario—primary infection with SARS-CoV-2 followed by IAV infection—SARS-CoV-2 replication was not significantly affected by the secondary IAV challenge (**Fig. 3G**). However, IAV replication was markedly impaired, with a 64% reduction in genome copy numbers compared with single IAV infection at 48 hpci (green bar, **Fig. 3H**). This effect was also supported and clearly visible by confocal microscopy (**Fig. 2**). This sequential coinfection (SARS-CoV-2 followed by IAV) did not significantly alter epithelial integrity relative to single infections, but it led to a pronounced increase in IL-6 and IFN-λ1 secretion compared to either single infection (green bars, **Fig. 3I**). The strong correlation between GCi and IFN-λ1 observed during single infections (**Fig. 1G**) was lost in this sequential coinfection context (**Fig. 3J**).

**Figure 3.**
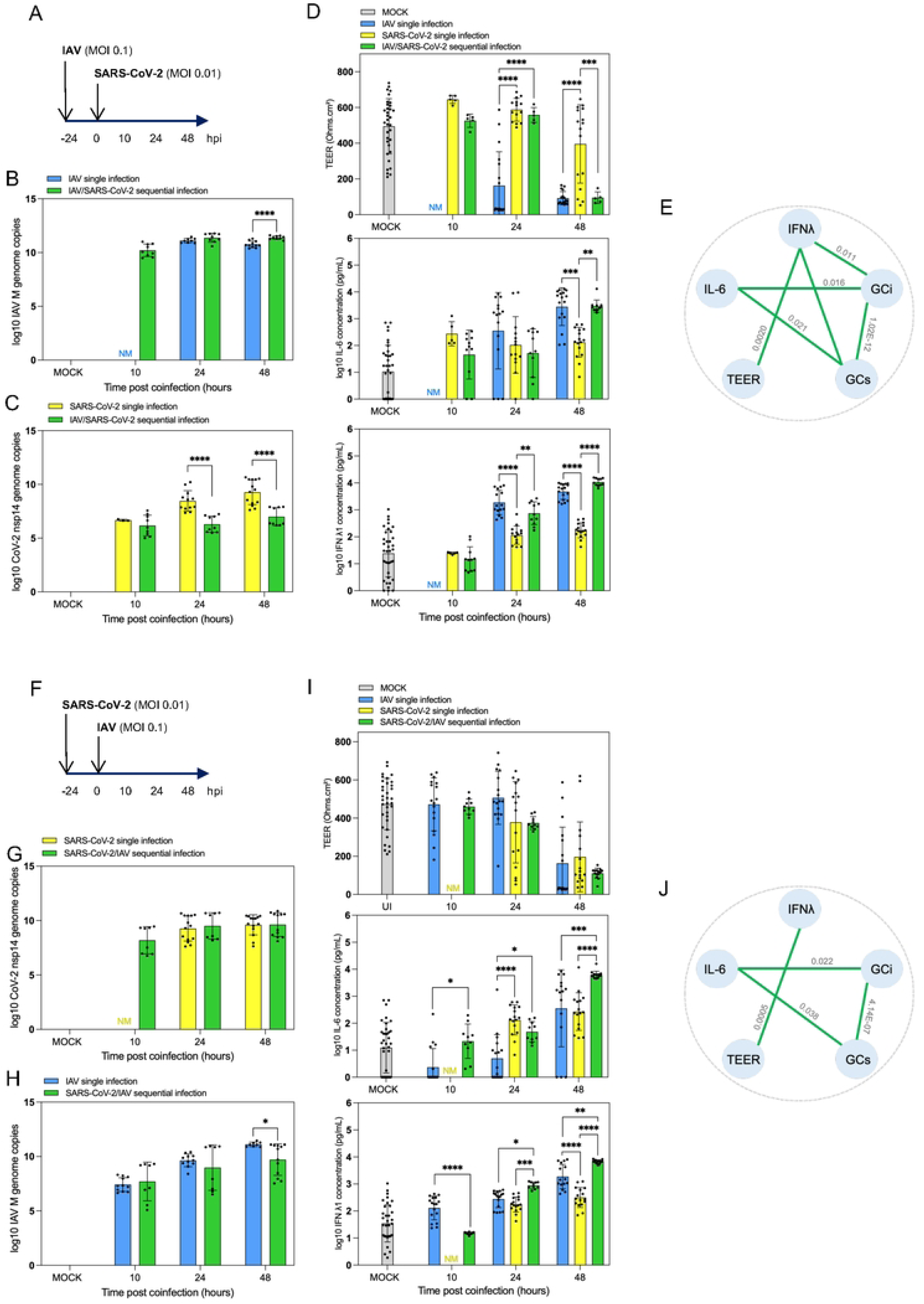
Sequential IAV/SARS-CoV-2 and SARS-CoV-2/IAV /IAV coinfections. (**A,F**) Reconstituted human bronchial epithelial tissues from six independent donors were infected or coinfected with IAV and/or SARS-CoV-2 at MOIs of 0.1 and 0.01, respectively, with a 24-hour delay between infections. Viral replication was monitored by RT-qPCR quantification of the IAV *M* gene (**B, G**) and the SARS-CoV-2 *nsp14* gene (**C, H**). Host parameters were assessed in parallel, including (**D, I**) TEER (Ω·cm²) as a measure of epithelial integrity, and secretion of IL-6 and IFN-λ1 (ELISA) to evaluate inflammatory and antiviral responses. (**E, J**) Partial correlation analysis was performed between viral and host parameters. Statistical analyses were conducted using one-way ANOVA followed by Dunnett’s, Sidak’s, and Tukey’s multiple-comparison tests. *, **, ***, and **** indicate adjusted p-values of 0.05, 0.01, 0.001, and 0.0001, respectively.

To further explore the mechanisms underlying IAV and SARS-CoV-2 interactions, we next analyzed the transcriptomic profiles of two sequential coinfection scenarios, comparing them to their corresponding single infection signatures at 48 h post-infection (i.e., IAV/SARS-CoV-2 versus SARS-CoV-2, and SARS-CoV-2/IAV versus IAV) (**Fig. 4A**). Differential expression analysis relative to the MOCK control revealed distinct transcriptomic signatures in both scenarios, characterized by large numbers of specific DEGs (n = 3,270 and 2,627 for IAV/SARS-CoV-2 and SARS-CoV-2/IAV, respectively) (**Fig. 4A**). This specificity largely reflected the impact of the primary infection (data not shown). Functional enrichment analysis identified highly similar GO terms between the two sequential coinfection conditions (**Fig. 4B** and **4C**), notably biological processes associated with the response to infection and inflammation. These findings are consistent with the markedly elevated IL-6 secretion observed during coinfection, particularly at 72 h post-coinfection (**Fig. 3D** and **3I**). We then analyzed the expression of the same panel of ISGs that had been previously examined under simultaneous coinfection (**Fig. 1K**). With the notable exception of IRF9, all ISGs were more strongly upregulated in coinfected samples compared to single infections, with expression levels more similar to those induced by IAV than by SARS-CoV-2 (**Fig. 4D**). These differential expression patterns likely contribute to the more pronounced effect of coinfection on SARS-CoV-2 replication compared to IAV in our experimental system.

**Figure 4.**
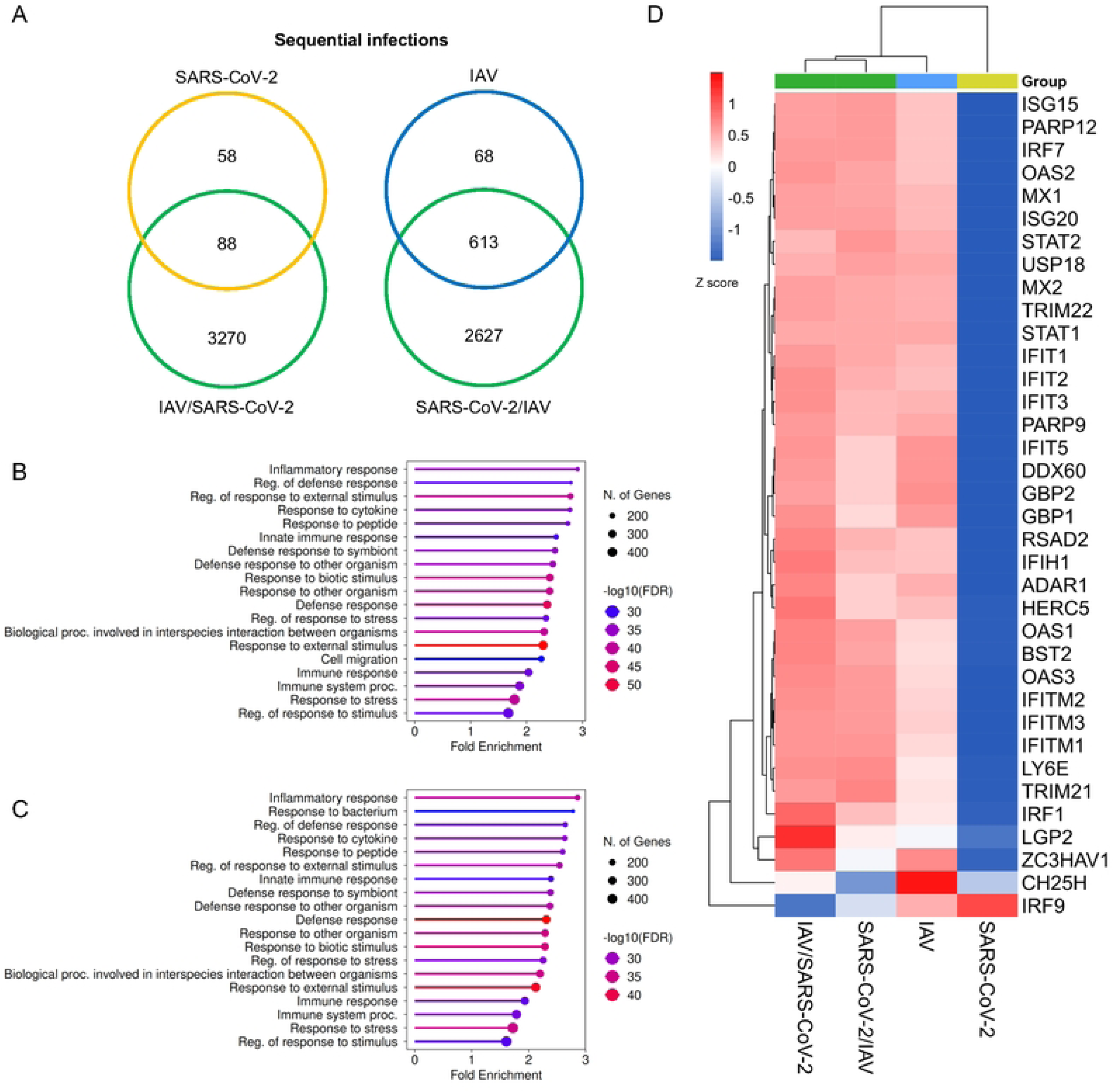
Transcriptomic profiling of sequential IAV/SARS-CoV-2 and SARS-CoV-2/IAV coinfections. Transcriptomic profiling was performed at 48 hpi using 3′ RNA-seq, selecting for each condition one representative infection from a single donor. (**A**) Differential expression analysis identified deregulated genes (DEGs) that were either shared or specific to each experimental condition. Functional enrichment of GO terms (Biological Processes) was performed using (**B**) the global or (**C**) the coinfection-specific upregulated gene signatures in IAV/SARS-CoV-2 or SARS-CoV-2/IAV coinfections. (**D**) Expression profiles of 36 ISGs known to participate in IAV and SARS-CoV-2 infection were analyzed, representing key innate immune pathways. Data are shown as heatmaps and expressed as Z-scores.

Altogether, these findings confirm the complex interplay between IAV and SARS-CoV-2 initially observed during simultaneous coinfections. However, they also reveal a key difference: the virus initiating the primary infection tends to gain a replicative advantage, or at least remains unaffected, by the secondary infection, in the case of IAV or SARS-CoV-2, respectively. These data highlight the critical importance of infection sequence in shaping both the magnitude and the nature of host responses, with direct consequences for the outcome of viral interactions.

### Simultaneous infection of HAE by IAV and RSV

Using the same experimental approach, we next examined the interaction between IAV and another major respiratory pathogen, respiratory syncytial virus (RSV), starting with the simultaneous coinfection scenario (**Fig. 5A**). Single RSV infection resulted in robust viral replication detectable as early as 10 hpi (red bars, **Fig. 5C**), with no significant impact on epithelial integrity (**Fig. 5D**). RSV infection also induced progressive secretion of IL-6 and IFN-λ1 throughout the infection course, reaching levels comparable to those measured during single IAV infection (**Fig. 5E** and **5F**). Correlation analysis between viral and host parameters identified significant associations between TEER and the cytokines IFN-λ1 and IL-6 (**Fig. 5G**). Under simultaneous coinfection conditions, we observed a transient but marked enhancement of IAV replication at 10 hpi (increase of 417%, purple bar, **Fig. 5B**), together with a pronounced inhibitory effect on RSV replication by at least 80% at all time points post-infection (purple bars, **Fig. 5C**). This strong negative impact of IAV on RSV was corroborated by confocal fluorescence microscopy, which showed a dramatic reduction in RSV-specific staining in co-infected samples (**Fig. 6**). In terms of host response, compared with single infections, the simultaneous coinfection did not significantly affect epithelial integrity, nor did it modify IL-6 or IFN-λ1 secretion levels beyond 24 hpi (**Fig. 5D** to **5F**). Notably, during coinfection, viral genome copy numbers for IAV (GCi) and RSV (GCr) were strongly correlated with each other and also showed a high correlation with IFN-λ1 levels, in contrast to what was observed during single infections (**Fig. 5G**).

**Figure 5.**
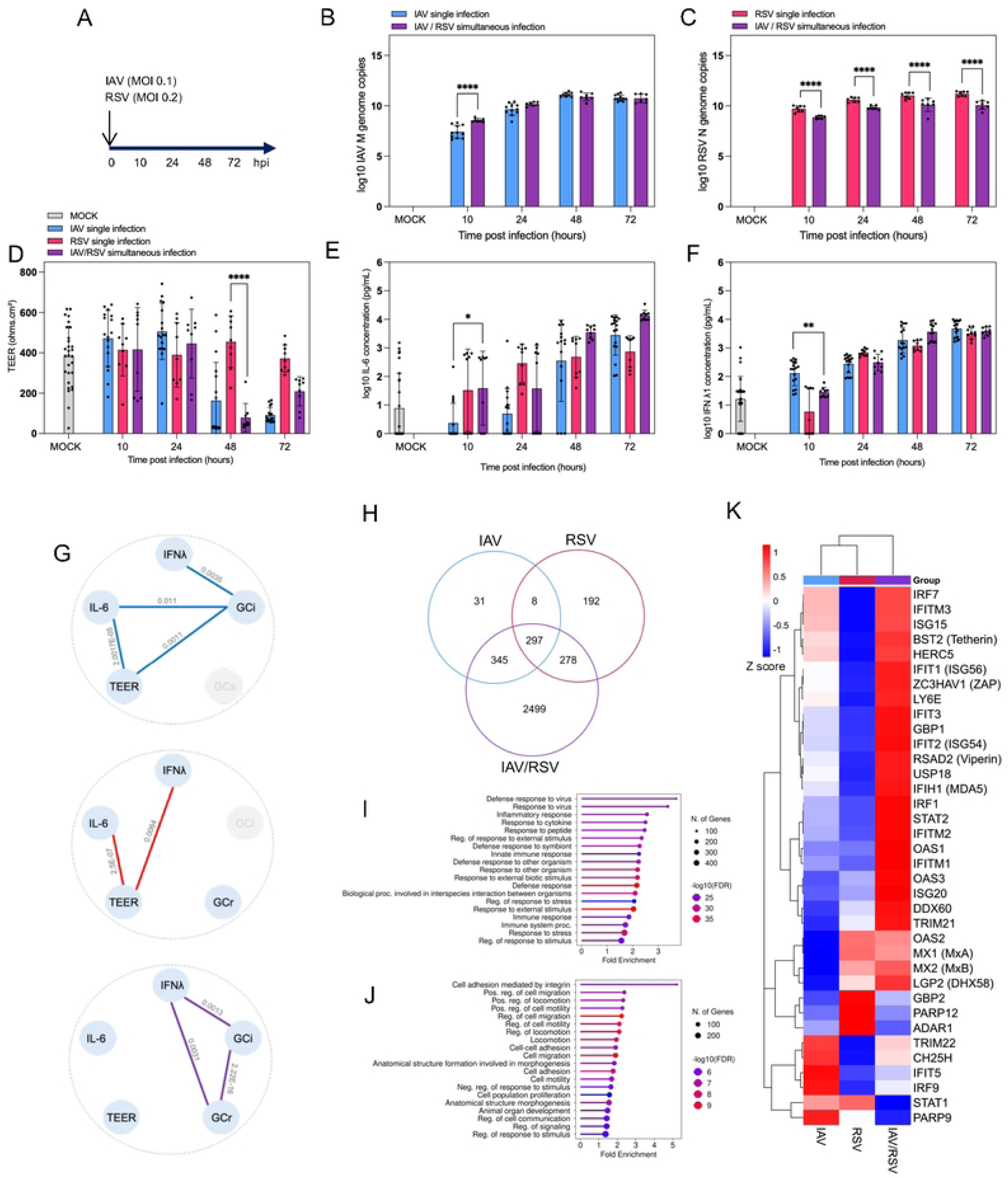
Simultaneous coinfection of IAV and RSV. (**A**) Reconstituted human bronchial epithelial tissues derived from six independent donors were infected or coinfected with IAV and/or RSV at MOIs of 0.1 and 0.2, respectively. Viral replication was monitored over time by quantifying genome copies of the IAV *M* gene (**B**) and the RSV *N* gene (**C**) using RT-qPCR. Host parameters were evaluated in parallel, including (**D**) TEER (Ω·cm²) as a proxy for epithelial integrity, and (**E–F**) secretion of IL-6 and IFN-λ1 (ELISA) to assess inflammatory and antiviral responses. (**G**) Partial correlation analysis was performed between viral and host parameters. To further characterize host transcriptional responses, transcriptomic profiling was performed at 48 hpi using 3′ RNA-seq (**H–K**), selecting one representative infection per condition. (**H**) Differential expression analysis identified DEGs either shared or specific among conditions. (**I–J**) Functional enrichment analysis of GO terms (Biological Processes) was conducted for global and coinfection-specific upregulated gene signatures. (**K**) Expression profiles of 36 ISGs implicated in IAV and RSV infection were analyzed, representing major innate immune pathways. Data are shown as heatmaps and expressed as Z-scores. Statistical analyses for panels (**B–F**) were performed using one-way ANOVA with Dunnett’s, Sidak’s, and Tukey’s multiple-comparison tests. *, **, ***, and **** denote adjusted p-values of 0.05, 0.01, 0.001, and 0.0001, respectively.

**Figure 6.**
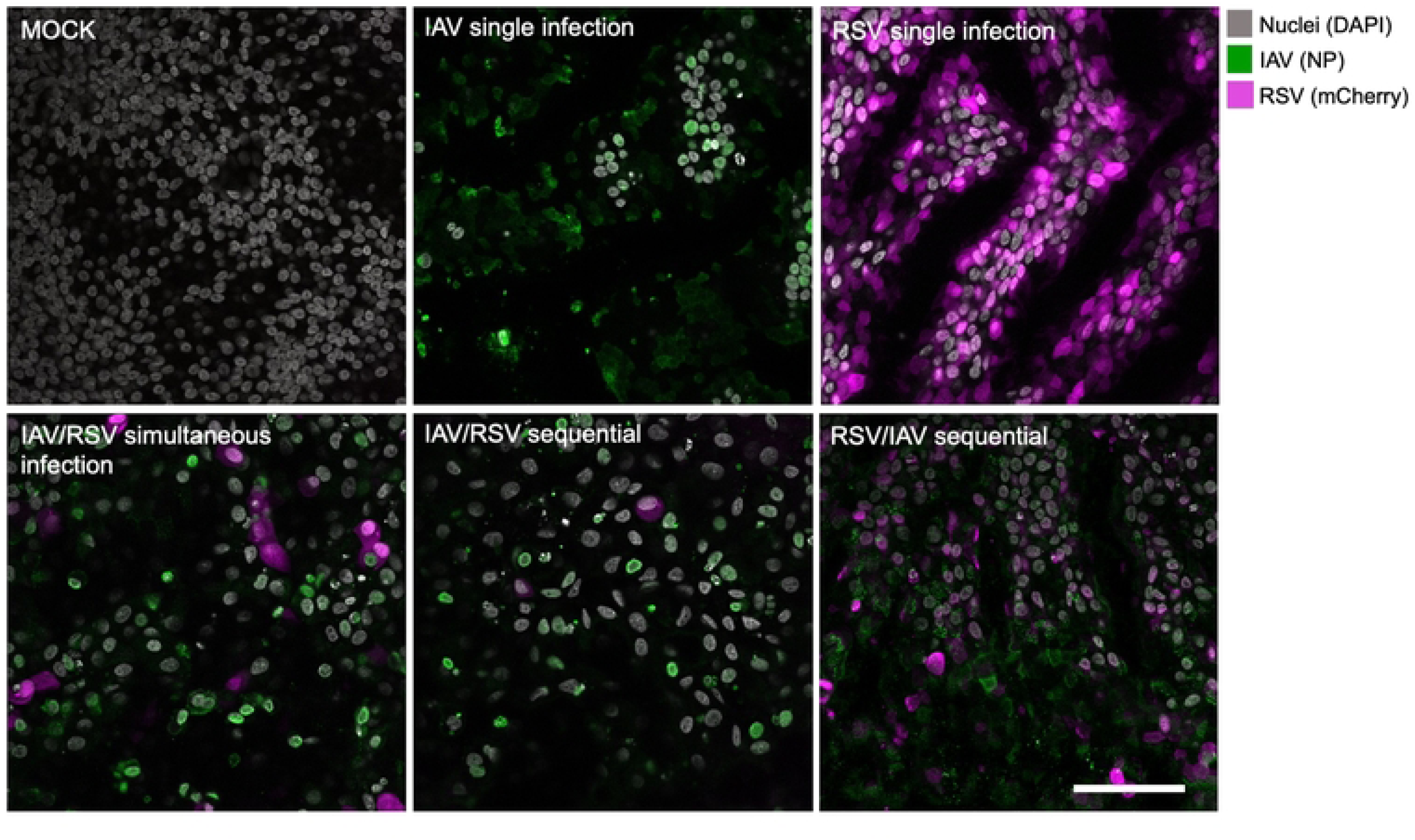
Confocal microscopy analysis of IAV and RSV coinfection scenarios. Representative confocal images of single infections or coinfections (simultaneous or sequential) were acquired at 48 hpi or 48 hpci, respectively. Confocal imaging was performed in oil immersion using a Leica SP5X microscope equipped with a 63× objective lens, and the images were processed with ImageJ Fiji software. Nuclei are shown in gray, IAV nucleoprotein (NP) in green, and RSV mCherry in magenta. Scale bar: 50 µm.

Differential gene expression analysis relative to the MOCK control (log₂ FC ≤ –1.5 or ≥ 1.5, adjusted p-value ≤ 0.05) identified a transcriptomic signature for simultaneous IAV/RSV coinfection comprising 3419 DEGs, with only 18.7% and 16.8% overlap with the signatures from IAV and RSV single infections, respectively (**Fig. 5H**). Remarkably, a significant fraction (n= 2,499, 73.13%) of this signature was specific to simultaneous coinfection (**Fig. 5H**). Functional enrichment analysis of upregulated genes revealed significant GO term enrichment in antiviral and immune response pathways (**Fig. 5I, S1 Fig. B**), similar to those identified in IAV and SARS-CoV-2 infections (**Supp. Fig. 1A**). In contrast, genes specifically upregulated during IAV/RSV coinfection were enriched in GO terms related to cell motility and adhesion (**Fig. 5J**), consistent with the marked epithelial morphological alterations observed by confocal microscopy between RSV single infections and IAV/RSV coinfections (**Fig. 6**). Comparison of ISG expression profiles across the three experimental conditions revealed overall higher expression in coinfection relative to single infections for many ISGs (e.g., IRF7, IFITM2, OAS3, MX1, MX2) (**Fig. 5K**). However, subsets of ISGs were preferentially upregulated in specific single infections—such as TRIM22, IFIT5, and IRF9 in IAV, and GBP2, PARP12, and ADAR1 in RSV (**Fig. 5K**).

Together, these data indicate that during simultaneous coinfection, IAV transiently benefits from the presence of RSV, while exerting a persistent and dominant suppressive effect on RSV replication. Despite this strong viral interference, host epithelial and cytokine responses remain largely comparable to those triggered by IAV alone, suggesting that IAV-driven host responses may overshadow or mask RSV-induced effects under these experimental conditions. Transcriptomic results indicate qualitative differences in the host innate immune response that may underlie the differential effects of coinfection on IAV and RSV replication under our experimental conditions. More broadly, consistent with our observations for IAV/SARS-CoV-2, the host response to coinfection appears to represent an integrated, non-additive interaction between the two viral infections.

### Sequential infections between IAV and RSV

We next investigated sequential coinfections between IAV and RSV, beginning with the scenario of a primary IAV infection followed by RSV infection 24 hours later (**Fig. 7A**). Under these experimental conditions, we observed a strong mutual negative interaction between the two viruses. Both exhibited markedly reduced replication levels in co-infected samples compared to single infections—by more than 70% for IAV as early as 24 hpi, and by more than 80% for RSV at 10 hpi (purple bars, **Fig. 7B** and **7C**). This mutual inhibition was also clearly apparent by confocal fluorescence microscopy, which revealed substantially weaker viral staining signals at 48 hpi in co-infected samples (**Fig. 6**). Regarding host parameters, the overall effects of sequential coinfection remained similar to those observed during single IAV infection (purple bars, **Fig. 7D**). Interestingly, in contrast to what was observed during simultaneous coinfection, correlations between viral genome copies (GCi and GCr) and IFN-λ1 levels were lost; instead, both GCi and GCr correlated with IL-6 secretion. The two viral replication parameters (GCi and GCr) remained strongly correlated with each other (**Fig. 7E**). Transcriptomic profiling of the host response further supported these observations, reflecting a global transcriptional signature consistent with strong reciprocal viral interference.

**Figure 7.**
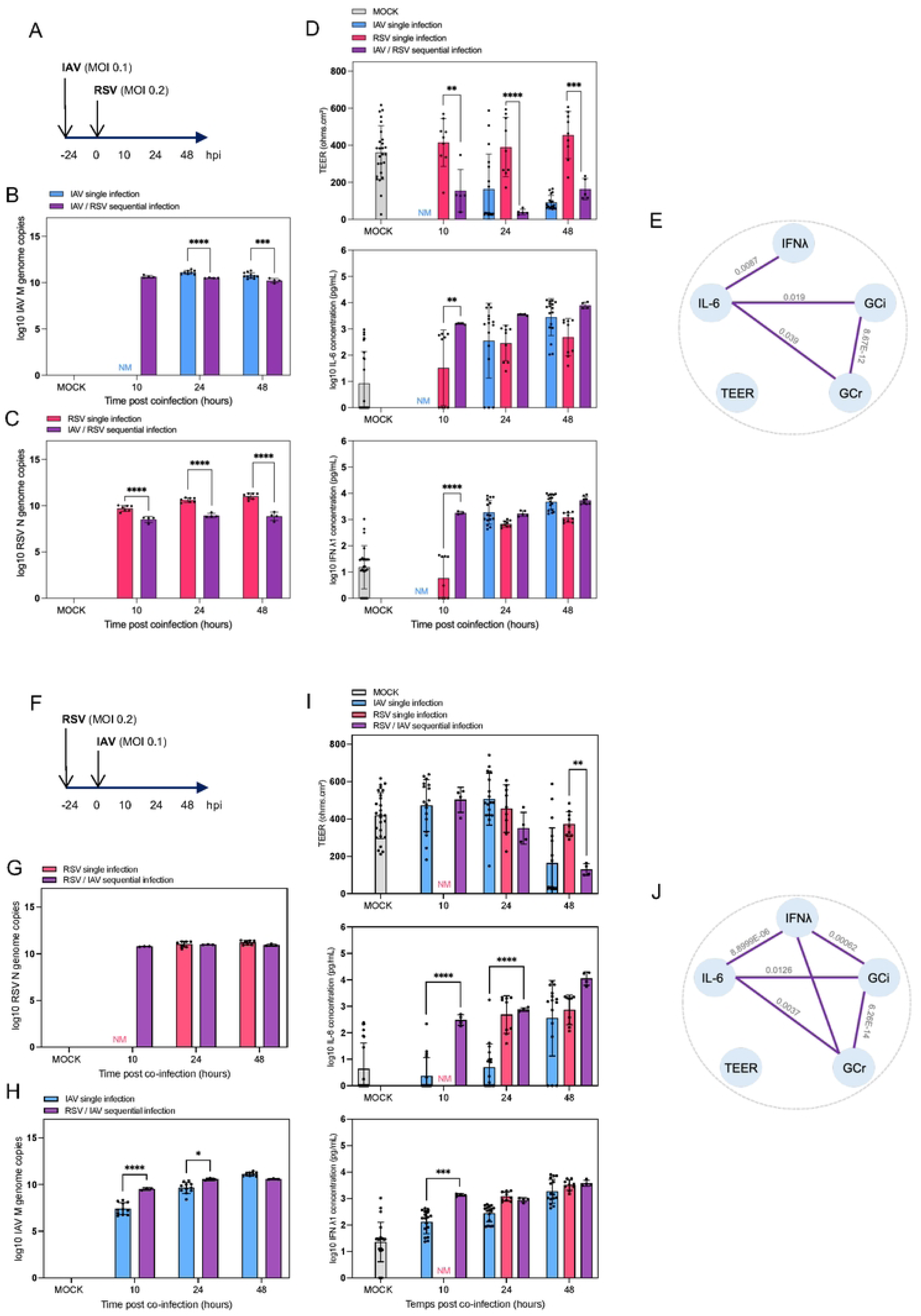
Sequential IAV/RSV and RSV/IAV coinfections. (**A, F**) Reconstituted human bronchial epithelial tissues from two independent donors were infected or coinfected with IAV and/or RSV at MOIs of 0.1 and 0.2, respectively, with a 24-hour delay between infections. Viral replication was quantified by RT-qPCR targeting the IAV *M* gene (**B, G**) and the RSV *N* gene (**C, H**). (**D, I**) TEER (Ω·cm²) was measured to assess epithelial integrity, and IL-6 and IFN-λ1 secretion was quantified by ELISA to monitor inflammatory and antiviral responses. (**E, J**) Partial correlation analysis was performed between viral and host parameters. Statistical analyses were conducted using one-way ANOVA with Dunnett’s, Sidak’s, and Tukey’s multiple-comparison tests. *, **, ***, and **** indicate adjusted p-values of 0.05, 0.01, 0.001, and 0.0001, respectively.

In the opposite sequential scenario—primary RSV infection followed by IAV infection (**Fig. 7F**); the pattern of viral interaction differed markedly. RSV replication was not significantly affected by subsequent IAV infection (**Fig. 7G**), whereas IAV replication was dramatically enhanced, increasing more than 46-fold compared to single IAV infection at 10 hpi (**Fig. 7H**). This boost in IAV replication was clearly evident by confocal microscopy (**Fig. 6**), which showed strong IAV staining overlapping with RSV-positive foci, likely corresponding to RSV-induced syncytia (data not shown). This sequential coinfection (RSV followed by IAV) produced an impact on epithelial integrity comparable to that observed during single IAV infection, but notably resulted in a pronounced increase in IL-6 and IFN-λ1 secretion at 10 and 24 hpci (purple bars, **Fig. 7I**). Correlation analysis among viral and host parameters indicated strong associations between all variables except TEER (**Fig. 7J**).

To gain deeper insight into the mechanisms governing IAV and RSV interactions, we analyzed transcriptomic profiles from two sequential coinfection scenarios, comparing them to their respective single infection signatures at 48 h post-infection (i.e., IAV/RSV versus RSV, and RSV/IAV versus IAV) (**Fig. 8A**). Differential expression analysis relative to the MOCK control revealed distinct transcriptomic signatures for both sequential coinfection conditions, with numerous specific DEGs (n = 2,161 and 2,038 for IAV/RSV and RSV/IAV, respectively) (**Fig. 8A**). This specificity largely reflected the influence of the primary infection (data not shown). Functional enrichment analysis identified highly similar GO terms between the IAV/RSV (**Fig. 8B**) and RSV/IAV (**Fig. 8C**) conditions, primarily involving pathways related to the response to infection. Examination of ISG expression patterns revealed distinct regulatory profiles across experimental conditions. Several ISGs displayed higher expression in both sequential coinfections (e.g., IFITM1, IFITM2, OAS1, TRIM21, IFIT1, ISG15, RSAD2, BST2) compared with single infections, while others were more specifically induced during RSV/IAV coinfection (e.g., IRF1, USP18, IFIT3, IFIT2, IFITM3) (**Fig. 8D**). These findings, similar to those observed for simultaneous coinfection, point to qualitative differences in the host innate immune response that contribute to the differential impact of coinfection on IAV and RSV replication under our experimental conditions.

**Figure 8.**
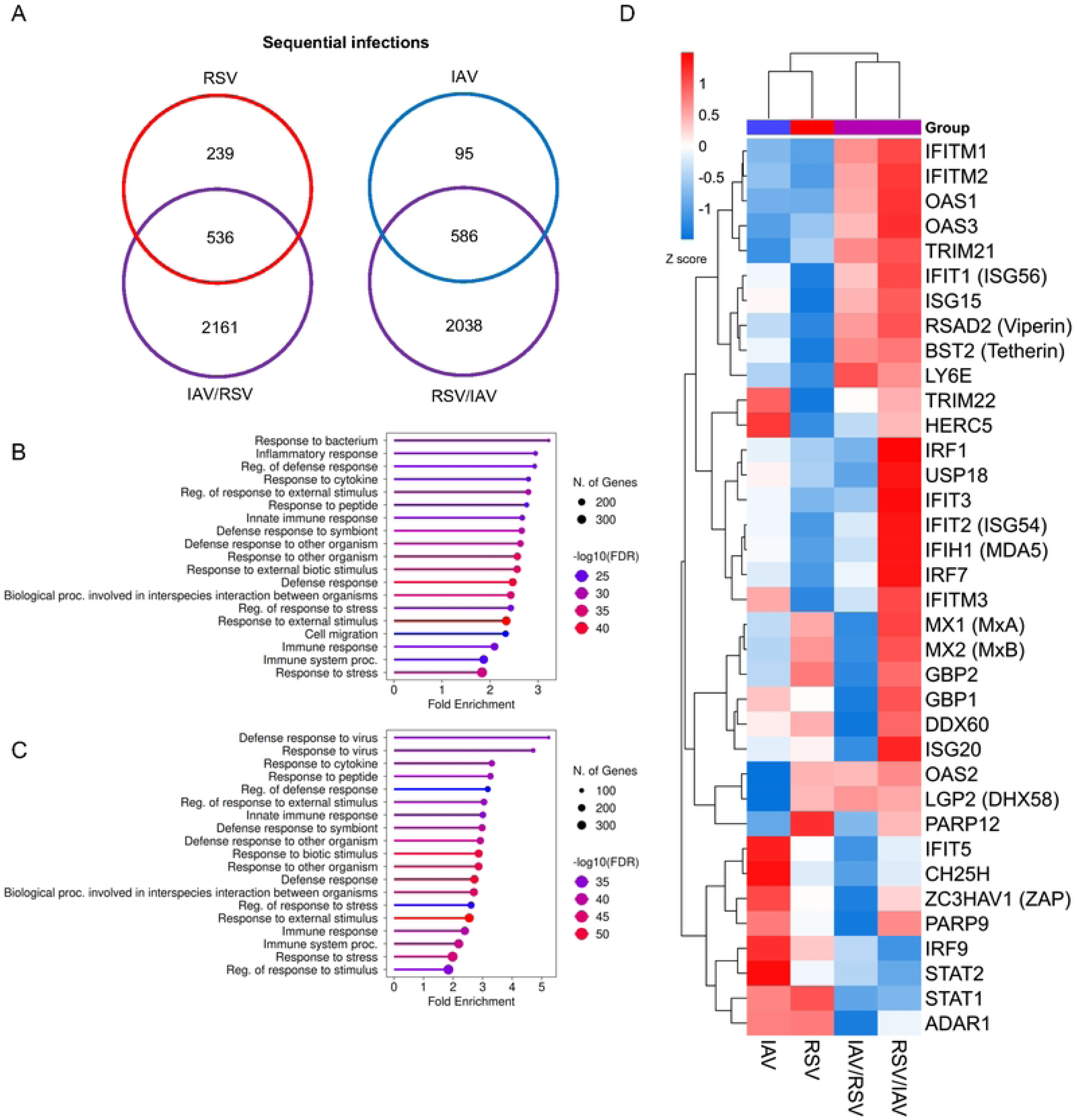
Transcriptomic profiling of sequential IAV/RSV and RSV/IAV coinfections. Transcriptomic profiling was performed at 48 hpi using 3′ RNA-seq, selecting one representative infection per condition. (**A**) Differential expression analysis identified DEGs that were either shared or specific among experimental conditions. Functional enrichment analysis of GO terms (Biological Processes) was performed for (**B**) global and (**C**) coinfection-specific upregulated gene signatures in IAV/RSV or RSV/IAV coinfections. (**D**) Expression profiles of 36 ISGs known to participate in IAV and RSV infection were analyzed, representing key innate immune pathways. Data are shown as heatmaps and expressed as Z-scores.

Altogether, these results highlight the complex and asymmetric nature of interactions between IAV and RSV during sequential infections. The directionality of infection plays a crucial role in determining the outcome of viral replication and host responses: while IAV tends to dominate and suppress RSV in most contexts, pre-existing RSV infection can, conversely, strongly enhance IAV replication, potentially through RSV-induced cellular remodeling or syncytia formation that facilitates IAV spread.

## DISCUSSION

The emergence of a new respiratory virus during the COVID-19 pandemic, alongside the multiple co-circulating respiratory viruses already known, has renewed interest in understanding their interactions and the consequences for disease severity and clinical management. Interactions between respiratory viruses are complex and can be antagonistic (viral interference), synergistic, or neutral, with implications at multiple levels that affect viral replication, pathogenicity, and epidemiological dynamics. In vitro studies—particularly those using reconstituted human airway epithelium (HAE) models—are essential for deciphering these mechanisms at the cellular level. In this study, we focused on two clinically relevant combinations of respiratory viruses [22,23], both involving influenza A virus (IAV, H1N1 subtype), either with SARS-CoV-2 or respiratory syncytial virus (RSV). Our aim was to systematically evaluate different coinfection scenarios in a human bronchial epithelium model over time, monitoring both viral replication and the host response in parallel.

Our findings regarding interactions between IAV and SARS-CoV-2 revealed a complex and asymmetric relationship. The primary infection generally maintained or gained a replicative advantage, while replication of the secondary virus was negatively impacted—consistent with our previous observations [21]. However, our data indicate that IAV exerts a much more potent inhibitory effect on SARS-CoV-2 than vice versa (**Fig. 3**), likely due to a more pronounced immune and inflammatory response induced by IAV infection, as evidenced by higher IL-6 and IFN-λ levels and strong upregulation of ISGs. These results suggest a unidirectional interference favoring IAV, in agreement with several prior reports. For example, Dee et al. showed that IAV (H3N2) inhibited replication of SARS-CoV-2 (Beta variant) in differentiated human bronchial epithelial cells (HBECs), with IAV triggering a much stronger IFN response [24]. Similarly, Cheemarla et al. demonstrated that IAV (H1N1pdm09) elicited a more robust interferon response than SARS-CoV-2 (WA/01 strain). In their sequential infection model (IAV three days before SARS-CoV-2), IAV suppressed SARS-CoV-2 replication by more than 4 log₁₀ at 72 h post-infection, whereas SARS-CoV-2 had no impact on IAV [25].

More recently, Gilbert-Girard et al. confirmed, in reconstituted nasal epithelium, a marked inhibitory effect of IAV (H1N1 and H3N2) on SARS-CoV-2 (D614 and Omicron variants), but not vice-versa, across both simultaneous and sequential infection scenarios [26]. In line with our findings, they attributed this unidirectional interference to higher IFN-λ production and stronger ISG upregulation following IAV infection. These in vitro results are further supported by in vivo observations: Zhang et al. reported that prior IAV (H1N1) infection in Syrian hamsters reduced SARS-CoV-2 lung viral load more effectively than the reverse sequence, although both coinfection conditions enhanced tissue damage [27]. This additive effect on epithelial damage parallels observations of compromised epithelial integrity [21].

In contrast to the strictly unidirectional interference seen in most reports, our study also detected bidirectional negative interference, particularly in simultaneous coinfection from 48 hpi onward (**Fig. 1**). Similar mutual interference has been described between IAV (H1N1pdm09) and SARS-CoV-2 (D614G variant), with >2 log₁₀ reductions in replication of both viruses under sequential infection conditions [28], as well as between influenza B virus and SARS-CoV-2 (Omicron variant) [29]. By contrast, Essaidi-Laziosi et al. reported that primary IAV (H1N1pdm09) infection reduced secondary SARS-CoV-2 replication, whereas primary SARS-CoV-2 infection had no effect on IAV replication [15].

Some studies have even suggested pro-viral effects in IAV/SARS-CoV-2 coinfections, consistent with our transient early replication “boost” observed for IAV (**Fig. 1B** and **Fig. 3B**). Bai et al. showed that prior influenza A infection enhanced SARS-CoV-2 infectivity in several cell lines (Vero E6, A549, Calu-3) and in K18-hACE2 transgenic mice infected 12 hours apart [30]. Conversely, when SARS-CoV-2 preceded IAV (H1N1pdm09), SARS-CoV-2 dominated replication, with higher viral loads, histopathological scores, and cytokine/chemokine expression levels than in single infections [27]. Collectively, these data, including ours, illustrate that IAV/SARS-CoV-2 interactions extend beyond simple negative interference, with differences likely arising from virus strain variability and divergent experimental designs. In sequential infections, the timing between inoculations appears crucial, potentially leading to opposite outcomes—consistent with discrepancies between this study and our previous work, which used a 48-hour interval instead of a 24-hour interval [21].

Interactions between IAV and RSV were also found to be complex and asymmetric. Overall, IAV tended to gain a replicative advantage over RSV in most contexts. Few studies have explored IAV–RSV interactions, but Gilbert-Girard et al. recently reported that, in bronchiolar HAE models, sequential IAV/RSV infections (4-day interval) resulted in a marked reduction of RSV viral load, whereas the reverse sequence did not affect IAV [26]. Similarly, in MDCK cells, IAV infection reduced RSV replication, with the magnitude of interference depending on the timing of infection [31] (Sinjoh et al., 2000). In other coinfection contexts (e.g., SARS-CoV-2 or rhinovirus), RSV generally appeared more susceptible to interference [28,32].

Interestingly, our data reveal that prior RSV infection can substantially enhance IAV replication, an observation not previously described in the literature (**Fig. 5B**). One possible explanation is that RSV-induced syncytia formation may facilitate the local spread of IAV within the epithelium. A similar mechanism has been reported for hPIV-2 and IAV coinfection in Vero cells [33], and the observation that dual-infected cells can produce hybrid viral particles supports this hypothesis [20].

Regarding the mechanisms underlying interactions between IAV and other viruses such as SARS-CoV-2 or RSV, our data and prior studies converge on the pivotal role of the interferon (IFN) response in determining coinfection outcomes. The magnitude of IFN induction and the capacity of each virus to counteract or evade it, both highly virus- and strain-dependent factors are key determinants and complicate mechanistic dissection at the cellular level. Several studies have employed IFN-pathway inhibitors, such as BX95 (a PDK1 inhibitor) or ruxolitinib (a JAK inhibitor) to demonstrate the central impact of IFN signaling on coinfection dynamics [24,26,28]. Reconstituted human respiratory epithelium models of nasal, bronchiolar, or bronchial origin—including ours—consistently point to IFN-λ, known to play a dominant role at epithelial surfaces [34], as a key mediator of virus–virus interactions [26,28].

It is increasingly recognized that viral interactions are not limited to IFN-mediated interference; numerous other mechanisms contribute to either negative or positive interactions among respiratory viruses [6]. Our observations of positive virus–virus interactions exemplify this complexity. Transcriptional profiling in our study revealed that while global coinfection signatures (IAV/RSV and IAV/SARS-CoV-2) were strongly enriched for immune response genes, coinfection-specific signatures highlighted distinct cellular pathways. Specifically, IAV/SARS-CoV-2 coinfection showed enrichment in metabolic pathways, including glycolysis, and in apoptosis-related pathways (**Fig. 1J**), consistent with established roles of these processes in IAV and SARS-CoV-2 infection [35,36]. The specific activation of these pathways during coinfection suggests additive and/or synergistic effects that may contribute to the greater epithelial damage observed, as evidenced by sharp declines in transepithelial electrical resistance (e.g., **Fig. 1D**) and the presence of apoptotic cells under electron microscopy [21]. Similarly, the IAV/RSV-specific signature showed significant enrichment in genes involved in cell motility and adhesion (**Fig. 5J**). These findings suggest that IAV/RSV coinfection may amplify the well-documented effect of RSV on adhesion molecule expression (e.g., ICAM-1 and VCAM-1), which are thought to play important roles in RSV pathophysiology [37]. This is consistent with our confocal microscopy observations showing marked epithelial morphological changes following RSV or IAV/RSV infection (**Fig. 6**).

Overall, our host-response analyses demonstrate that the response to coinfection is far from a simple sum of single-virus responses; rather, it exhibits specific, emergent properties. Characterizing these unique transcriptional signatures may help elucidate the potential mechanisms underlying clinical aggravation during coinfection, as suggested by both in vitro and clinical studies [7,38,39].

Despite the strong physiological relevance of reconstituted human airway epithelium models used in this study and in many others, these systems have limitations, particularly the absence of immune cells. The development of next-generation multicellular models incorporating components such as macrophages or dendritic cells [40] will undoubtedly advance our understanding of virus–virus interactions in the respiratory tract. Likewise, the advent of primary CRISPR–Cas9–modified airway epithelial models will enable a finer dissection of host factors, not only in immune pathways but also across other key cellular processes, such as metabolic or apoptotic pathways.

In conclusion, our study advances understanding of interactions between the influenza virus and SARS-CoV-2 or RSV, integrating longitudinal analysis of both viral and host parameters. Our results underscore the complexity of these interactions, which range from mutual to unidirectional interference depending on the viral models and highlight the central role of innate immune responses and the sequence of infection in determining coinfection outcomes. We believe that combining diverse experimental models, in vitro and in vivo, with mathematical modeling will be essential to fully elucidate the underlying mechanisms and capture the true complexity of virus–virus interactions, with implications for future clinical management, epidemic forecasting, and the development of antiviral strategies.

## MATERIALS AND METHODS

### Reconstituted Human Airway Epithelium (HAE)

MucilAir™ human airway epithelia (HAE), reconstituted from primary human bronchial cells obtained from biopsies, were purchased from Epithelix (MucilAir, Switzerland). Cultures were maintained at an air–liquid interface (ALI) in a proprietary medium using Costar Transwell inserts (Corning) according to the manufacturer’s instructions. To account for donor variability, bronchial HAE was derived from multiple donors differing in age, sex, and ethnicity (see **S2 Table** for donor information). Epithelial integrity was assessed by measuring transepithelial electrical resistance (TEER, Ω·cm²) using an EVOM2 Epithelial Volt/Ohm meter (World Precision Instruments). IL-6 and IFN-λ secretion were quantified in basal medium using commercial ELISA kits (Mabtech, ref. 3460-1H; Invitrogen, ref. 88-7296-22) according to the manufacturer’s protocols. The substrate 3,3′,5,5′-tetramethylbenzidine (TMB) was purchased from Cell Signaling Technology.

### Viruses and Infection of HAE

The SARS-CoV-2 strain used in this study was isolated from a patient enrolled in a French clinical cohort of COVID-19 patients (ClinicalTrials.gov identifier NCT04262921; [21]. The sequence is available in the GISAID EpiCoV™ database under the accession BetaCoV/France/IDF0571/2020 (EPI_ISL_411218). Viral stocks were amplified and titrated by determination of the 50% tissue culture infectious dose (TCID₅₀/mL) on Vero cells (ATCC CRL-1586). The influenza A/Lyon/969/2009 (H1N1) strain was propagated and titrated on MDCK cells (ATCC CCL-34), as described previously [41]. A recombinant RSV A2 strain expressing mCherry (RSV-mCherry, kindly provided by Jean-François Eléouët, INRAE, Jouy-en-Josas, France) was propagated and quantified on LLC-MK2 cells (ATCC CCL-7), as described previously [42]. For infection, HAE were gently washed twice at the apical surface with Opti-MEM (Life Technologies) and subsequently mock-infected (Opti-MEM) or infected with SARS-CoV-2, IAV, or RSV at multiplicities of infection (MOI) of 0.01, 0.1, and 0.2, respectively. At defined time points, apical washes and basal media were collected and stored at −80°C for further analyses. Cells were lysed for total RNA extraction, which was used for RT-qPCR and RNA-seq. All experiments involving infectious SARS-CoV-2 material were performed in a biosafety level 3 (BSL-3) facility, while infections involving only IAV and/or RSV were conducted in a biosafety level 2 (BSL-2) laboratory.

### Viral Quantification by RT-qPCR

Viral replication was quantified by measuring viral RNA levels. Briefly, HAE were lysed in RLT buffer (Qiagen RNeasy® Plus Mini Kit, ref. 74136) and scraped before collection. Total RNA was extracted using the same kit according to the manufacturer’s instructions. RNA quantification was performed using a one-step RT-qPCR (Luna Universal Probe qPCR Master Mix, New England Biolabs, ref. M3004) on an Applied Biosystems StepOnePlus Real-Time PCR system. Each sample was analyzed in duplicate. Primers and probes used in this study are listed in **S3 Table**. Threshold cycle (Cₜ) values were converted to viral genome copy numbers using standard curves generated from known RNA standards.

### Immunofluorescence Confocal Microscopy

Mock- and virus-infected, or coinfected, epithelia were fixed with 4% paraformaldehyde (PFA) for 10 min and permeabilized twice for 25 min with 0.1% Triton X-100 in PBS, applied to both the apical and basal surfaces. Blocking was performed with 1% fetal bovine serum (FBS) in PBS on both sides. The apical surface was incubated for 2 h with primary antibodies against viral proteins, followed by three 10-minute washes in PBS containing 0.1% Triton X-100. Cells were then incubated for 30 min with fluorophore-conjugated secondary antibodies, washed again three times, and stained with DAPI (1 µg/mL, Invitrogen, ref. 66248) for nuclear visualization. Epithelial monolayers were mounted in Fluoromount (Invitrogen, ref. 00-4958-02). Confocal images were acquired using a Leica TCS-SP5X microscope (Wetzlar, Germany) and processed with ImageJ FIJI software https://imagej.net/software/fiji/. The complete list of antibodies and corresponding references is provided in **Supplementary Table 3**.

### Transcriptomic Profiling

RNA library preparation was realized following the manufacturer’s recommendations (QuantSeq 3’ mRNA-Seq V2 Library Prep Kit from LEXOGEN). Final samples pooled library prep were sequenced on an ILLUMINA NextSeq 2000 with a P2-100 cycles cartridge (1*400 million 100-base reads), corresponding to 1*13 million reads per sample after demultiplexing. The RNA-seq data analysis was performed using BioJupies [43]. Raw counts were normalized to log10-Counts Per Million (logCPM) by dividing each column by the total sum of its counts, multiplying it by 106, followed by the application of a log10-transform. The gene expression signature was generated by comparing gene expression levels between the control group and the experimental group using the limma R package [44]. Functional gene-set enrichment was performed using ShinyGO 0.85 https://bioinformatics.sdstate.edu/go/ [45]. Heatmap visualization was performed using SRplot https://www.bioinformatics.com.cn/srplot [46]. All sequencing data have been deposited in the ArrayExpress collection in BioStudies (https://www.ebi.ac.uk/biostudies/arrayexpress) under accession reference E-MTAB-16265.

## Statistical Analyses

Statistical analyses were performed using one-way ANOVA followed by Dunnett’s, Sidak’s, or Tukey’s multiple comparison tests, as indicated. Partial correlation analyses were conducted to evaluate the relationship between two parameters while controlling for the influence of others [47].

## ACKNOWLEDGMENTS

We thank Clément Fage and all members of the VirPath-CIRI lab for the critical reading of the manuscript and helpful discussions. The authors would like to thank the Centre d’Imagerie Quantitative Lyon-Est (CIQLE), Université Claude Bernard Lyon 1, for support with confocal microscopy. This work benefited from equipment and services from the iGenSeq core facility (Genotyping and sequencing), at Paris Brain Institute (PBI).

## AUTHOR CONTRIBUTIONS

Conceptualization: AG, AP, OT; Methodology: AG, PL, ALG, JG, FG, AP, OT; Investigation: AG, CT, EP, PL, OT; Writing – original draft: AG, OT; Writing Review & Editing: AG, AP, OT; Funding acquisition: OT; Supervision: AP, OT.

## Supplementary information captions

**S1 Figure. Functional enrichment analyses for IAV/SARS-CoV-2 and IAV/RSV simultaneous coinfections.** Functional enrichment analysis of GO terms (Biological Processes) was conducted for global and coinfection-specific upregulated gene signatures in the context of IAV/SARS-CoV-2 (**A**) and IAV/RSV (**B**) simultaneous coinfections.

**S2 Table. List of HAE donors in this study**

**S3 Table. List of primers/probes used in this study.**

## REFERENCES

1. Safiri S, Mahmoodpoor A, Kolahi A-A, Nejadghaderi SA, Sullman MJM, Mansournia MA, et al. Global burden of lower respiratory infections during the last three decades. Front Public Health. 2022;10: 1028525. doi:10.3389/fpubh.2022.1028525

2. Kang L, Jing W, Liu J, Liu M. Trends of global and regional aetiologies, risk factors and mortality of lower respiratory infections from 1990 to 2019: An analysis for the Global Burden of Disease Study 2019. Respirology. 2023;28: 166–175. doi:10.1111/resp.14389

3. Jin X, Ren J, Li R, Gao Y, Zhang H, Li J, et al. Global burden of upper respiratory infections in 204 countries and territories, from 1990 to 2019. EClinicalMedicine. 2021;37: 100986. doi:10.1016/j.eclinm.2021.100986

4. Shi T, Arnott A, Semogas I, Falsey AR, Openshaw P, Wedzicha JA, et al. The Etiological Role of Common Respiratory Viruses in Acute Respiratory Infections in Older Adults: A Systematic Review and Meta-analysis. J Infect Dis. 2020;222: S563–S569. doi:10.1093/infdis/jiy662

5. Aberle JH, Aberle SW, Pracher E, Hutter H-P, Kundi M, Popow-Kraupp T. Single versus dual respiratory virus infections in hospitalized infants: impact on clinical course of disease and interferon-gamma response. Pediatr Infect Dis J. 2005;24: 605–610. doi:10.1097/01.inf.0000168741.59747.2d

6. Trepat K, Gibeaud A, Trouillet-Assant S, Terrier O. Exploring viral respiratory coinfections: Shedding light on pathogen interactions. PLoS Pathog. 2024;20: e1012556. doi:10.1371/journal.ppat.1012556

7. Stowe J, Tessier E, Zhao H, Guy R, Muller-Pebody B, Zambon M, et al. Interactions between SARS-CoV-2 and influenza, and the impact of coinfection on disease severity: a test-negative design. Int J Epidemiol. 2021;50: 1124–1133. doi:10.1093/ije/dyab081

8. Guan Z, Chen C, Li Y, Yan D, Zhang X, Jiang D, et al. Impact of Coinfection With SARS-CoV-2 and Influenza on Disease Severity: A Systematic Review and Meta-Analysis. Frontiers in Public Health. 2021;9. Available: https://www.frontiersin.org/articles/10.3389/fpubh.2021.773130

9. Lansbury L, Lim B, Baskaran V, Lim WS. Co-infections in people with COVID-19: a systematic review and meta-analysis. J Infect. 2020;81: 266–275. doi:10.1016/j.jinf.2020.05.046

10. Carstens G, Kozanli E, Bulsink K, McDonald SA, Elahi M, de Bakker J, et al. Co-infection dynamics of SARS-CoV-2 and respiratory viruses in the 2022/2023 respiratory season in the Netherlands. J Infect. 2025;90: 106474. doi:10.1016/j.jinf.2025.106474

11. Waterlow NR, Flasche S, Minter A, Eggo RM. Competition between RSV and influenza: Limits of modelling inference from surveillance data. Epidemics. 2021;35: 100460. doi:10.1016/j.epidem.2021.100460

12. Kramer SC, Pirikahu S, Casalegno J-S, Domenech de Cellès M. Characterizing the interactions between influenza and respiratory syncytial viruses and their implications for epidemic control. Nat Commun. 2024;15: 10066. doi:10.1038/s41467-024-53872-4

13. Piret J, Boivin G. Viral Interference between Respiratory Viruses. Emerg Infect Dis. 2022;28: 273–281. doi:10.3201/eid2802.211727

14. Dee K, Goldfarb DM, Haney J, Amat JAR, Herder V, Stewart M, et al. Human Rhinovirus Infection Blocks Severe Acute Respiratory Syndrome Coronavirus 2 Replication Within the Respiratory Epithelium: Implications for COVID-19 Epidemiology. J Infect Dis. 2021;224: 31–38. doi:10.1093/infdis/jiab147

15. Essaidi-Laziosi M, Alvarez C, Puhach O, Sattonnet-Roche P, Torriani G, Tapparel C, et al. Sequential infections with rhinovirus and influenza modulate the replicative capacity of SARS-CoV-2 in the upper respiratory tract. Emerg Microbes Infect. 2022;11: 412–423. doi:10.1080/22221751.2021.2021806

16. Gilbert-Girard S, Piret J, Rhéaume C, Carbonneau J, Goyette N, Couture C, et al. Influenza A virus interferes with respiratory syncytial virus in mice and reconstituted human airway epithelium. Microbiol Spectr. 2025;13: e0318724. doi:10.1128/spectrum.03187-24

17. Pinky L, DeAguero JR, Remien CH, Smith AM. How Interactions during Viral-Viral Coinfection Can Shape Infection Kinetics. Viruses. 2023;15: 1303. doi:10.3390/v15061303

18. Hartwig SM, Miller AM, Varga SM. Respiratory Syncytial Virus Provides Protection against a Subsequent Influenza A Virus Infection. J Immunol. 2022;208: 720–731. doi:10.4049/jimmunol.2000751

19. George JA, AlShamsi SH, Alhammadi MH, Alsuwaidi AR. Exacerbation of Influenza A Virus Disease Severity by Respiratory Syncytial Virus Co-Infection in a Mouse Model. Viruses. 2021;13: 1630. doi:10.3390/v13081630

20. Haney J, Vijayakrishnan S, Streetley J, Dee K, Goldfarb DM, Clarke M, et al. Coinfection by influenza A virus and respiratory syncytial virus produces hybrid virus particles. Nat Microbiol. 2022;7: 1879–1890. doi:10.1038/s41564-022-01242-5

21. Pizzorno A, Padey B, Dulière V, Mouton W, Oliva J, Laurent E, et al. Interactions Between Severe Acute Respiratory Syndrome Coronavirus 2 Replication and Major Respiratory Viruses in Human Nasal Epithelium. J Infect Dis. 2022;226: 2095–2104. doi:10.1093/infdis/jiac357

22. Swets MC, Russell CD, Harrison EM, Docherty AB, Lone N, Girvan M, et al. SARS-CoV-2 co-infection with influenza viruses, respiratory syncytial virus, or adenoviruses. Lancet. 2022;399: 1463–1464. doi:10.1016/S0140-6736(22)00383-X

23. Zhang Y, Zhao J, Zou X, Fan Y, Xiong Z, Li B, et al. Severity of influenza virus and respiratory syncytial virus coinfections in hospitalized adult patients. J Clin Virol. 2020;133: 104685. doi:10.1016/j.jcv.2020.104685

24. Dee K, Schultz V, Haney J, Bissett LA, Magill C, Murcia PR. Influenza A and Respiratory Syncytial Virus Trigger a Cellular Response That Blocks Severe Acute Respiratory Syndrome Virus 2 Infection in the Respiratory Tract. J Infect Dis. 2023;227: 1396–1406. doi:10.1093/infdis/jiac494

25. Cheemarla NR, Watkins TA, Mihaylova VT, Foxman EF. Viral Interference During Influenza A-SARS-CoV-2 Coinfection of the Human Airway Epithelium and Reversal by Oseltamivir. J Infect Dis. 2024;229: 1430–1434. doi:10.1093/infdis/jiad402

26. Gilbert-Girard S, Piret J, Carbonneau J, Hénaut M, Goyette N, Boivin G. Viral interference between severe acute respiratory syndrome coronavirus 2 and influenza A viruses. PLoS Pathog. 2024;20: e1012017. doi:10.1371/journal.ppat.1012017

27. Zhang AJ, Lee AC-Y, Chan JF-W, Liu F, Li C, Chen Y, et al. Coinfection by Severe Acute Respiratory Syndrome Coronavirus 2 and Influenza A(H1N1)pdm09 Virus Enhances the Severity of Pneumonia in Golden Syrian Hamsters. Clin Infect Dis. 2021;72: e978–e992. doi:10.1093/cid/ciaa1747

28. Fage C, Hénaut M, Carbonneau J, Piret J, Boivin G. Influenza A(H1N1)pdm09 Virus but Not Respiratory Syncytial Virus Interferes with SARS-CoV-2 Replication during Sequential Infections in Human Nasal Epithelial Cells. Viruses. 2022;14: 395. doi:10.3390/v14020395

29. Reuss D, Brown JC, Sukhova K, Furnon W, Cowton V, Patel AH, et al. Interference between SARS-CoV-2 and influenza B virus during coinfection is mediated by induction of specific interferon responses in the lung epithelium. Virology. 2025;608: 110556. doi:10.1016/j.virol.2025.110556

30. Bai L, Zhao Y, Dong J, Liang S, Guo M, Liu X, et al. Coinfection with influenza A virus enhances SARS-CoV-2 infectivity. Cell Res. 2021;31: 395–403. doi:10.1038/s41422-021-00473-1

31. Shinjoh M, Omoe K, Saito N, Matsuo N, Nerome K. In vitro growth profiles of respiratory syncytial virus in the presence of influenza virus. Acta Virol. 2000;44: 91–97.

32. Essaidi-Laziosi M, Geiser J, Huang S, Constant S, Kaiser L, Tapparel C. Interferon-Dependent and Respiratory Virus-Specific Interference in Dual Infections of Airway Epithelia. Sci Rep. 2020;10: 10246. doi:10.1038/s41598-020-66748-6

33. Goto H, Ihira H, Morishita K, Tsuchiya M, Ohta K, Yumine N, et al. Enhanced growth of influenza A virus by coinfection with human parainfluenza virus type 2. Medical Microbiology and Immunology. 2016;205: 209. doi:10.1007/s00430-015-0441-y

34. Wells AI, Coyne CB. Type III Interferons in Antiviral Defenses at Barrier Surfaces. Trends Immunol. 2018;39: 848–858. doi:10.1016/j.it.2018.08.008

35. Liu X, Chen L, Niu H, Chen Y, Chen P, Liu L, et al. The bittersweet link between glucose metabolism, cellular microenvironment and viral infection. Virulence. 2025;16: 2554302. doi:10.1080/21505594.2025.2554302

36. Shao Q, Liu T, Hu B, Chen L. Interplay between autophagy and apoptosis in human viral pathogenesis. Virus Res. 2025;359: 199611. doi:10.1016/j.virusres.2025.199611

37. Wang SZ, Hallsworth PG, Dowling KD, Alpers JH, Bowden JJ, Forsyth KD. Adhesion molecule expression on epithelial cells infected with respiratory syncytial virus. Eur Respir J. 2000;15: 358–366. doi:10.1034/j.1399-3003.2000.15b23.x

38. Achdout H, Vitner EB, Politi B, Melamed S, Yahalom-Ronen Y, Tamir H, et al. Increased lethality in influenza and SARS-CoV-2 coinfection is prevented by influenza immunity but not SARS-CoV-2 immunity. Nat Commun. 2021;12: 5819. doi:10.1038/s41467-021-26113-1

39. Pott H, J LeBlanc J, S ElSherif M, Hatchette TF, McNeil SA, Andrew MK, et al. Clinical features and outcomes of influenza and RSV coinfections: a report from Canadian immunization research network serious outcomes surveillance network. BMC Infect Dis. 2024;24: 147. doi:10.1186/s12879-024-09033-5

40. Gibeaud A, Pizzorno A, Terrier O. In vitro modeling of influenza infection in the respiratory epithelium: advanced cellular models to better understand complex host-virus interactions. Curr Opin Virol. 2025;70: 101452. doi:10.1016/j.coviro.2025.101452

41. Terrier O, Textoris J, Carron C, Marcel V, Bourdon J-C, Rosa-Calatrava M. Host microRNA molecular signatures associated with human H1N1 and H3N2 influenza A viruses reveal an unanticipated antiviral activity for miR-146a. J Gen Virol. 2013;94: 985–995. doi:10.1099/vir.0.049528-0

42. Ogonczyk-Makowska D, Brun P, Vacher C, Chupin C, Droillard C, Carbonneau J, et al. Mucosal bivalent live attenuated vaccine protects against human metapneumovirus and respiratory syncytial virus in mice. NPJ Vaccines. 2024;9: 111. doi:10.1038/s41541-024-00899-9

43. Torre D, Lachmann A, Ma’ayan A. BioJupies: Automated Generation of Interactive Notebooks for RNA-Seq Data Analysis in the Cloud. Cell Syst. 2018;7: 556–561.e3. doi:10.1016/j.cels.2018.10.007

44. Ritchie ME, Phipson B, Wu D, Hu Y, Law CW, Shi W, et al. limma powers differential expression analyses for RNA-sequencing and microarray studies. Nucleic Acids Res. 2015;43: e47. doi:10.1093/nar/gkv007

45. Ge SX, Jung D, Yao R. ShinyGO: a graphical gene-set enrichment tool for animals and plants. Bioinformatics. 2020;36: 2628–2629. doi:10.1093/bioinformatics/btz931

46. Tang D, Chen M, Huang X, Zhang G, Zeng L, Zhang G, et al. SRplot: A free online platform for data visualization and graphing. PLoS One. 2023;18: e0294236. doi:10.1371/journal.pone.0294236

47. Baba K, Shibata R, Sibuya M. Partial Correlation and Conditional Correlation as Measures of Conditional Independence. Australian & New Zealand Journal of Statistics. 2004;46: 657–664. doi:10.1111/j.1467-842X.2004.00360.x

